# The distribution of highly deleterious variants across human ancestry groups

**DOI:** 10.1101/2025.01.31.635988

**Authors:** Anastasia Stolyarova, Graham Coop, Molly Przeworski

**Affiliations:** Dept of Biological Sciences, Columbia University; Dept of Evolution and Ecology and Center for Population Biology, University of California, Davis; Dept of Systems Biology, Columbia University

## Abstract

A major focus of human genetics is to map severe disease mutations. Increasingly that goal is understood as requiring huge numbers of people to be sequenced from every broadly-defined genetic ancestry group, so as not to miss “ancestry-specific variants.” Here, we argue that this focus is unwarranted. We start with first principles considerations, based on models of mutation-drift-selection balance, which suggest highly pathogenic mutations should be at similarly low frequencies across ancestry groups. Severe disease mutations tend to be strongly deleterious, and thus evolutionarily young, and are kept at relatively constant frequency through recurrent mutation. Therefore, highly pathogenic alleles are shared identical by descent within extended families, not broad ancestry groups, and sequencing more people should yield similar numbers regardless of ancestry. We illustrate these points using gnomAD genetic ancestry groupings, and show that the classes of variants most likely to be highly pathogenic, notably sets of loss of function alleles at strongly constrained genes, conform well to these predictions. While there are many important reasons to diversify genomic research, strongly deleterious alleles will be found at comparable rates in people of all ancestries, and the information they provide about human biology is shared across ancestries.

## Main text

The identification of pathogenic variants helps diagnose patients and point to genes whose disruptions are associated with disease outcomes, providing an entry point to understand pathophysiology and develop new drug targets^1–5^. These goals have been transformed by DNA sequencing technologies, and millions of people have now had their genomes sequenced. In fact, sample sizes are approaching saturation mutagenesis: in the absence of deleterious effects, almost every single nucleotide site will be variable as sample sizes approach tens of millions of people^6–8^.

To date, this massive sequencing effort has disproportionately sampled people of recent European ancestry^9–11^. This over-representation reflects inequities of many kinds, notably in access to health care, and risks perpetuating others. Notably, genomic predictions of complex disease susceptibility are currently much less accurate for people of non-European ancestries^12,13^ (as well as for people in distinct environmental settings^14,15^). Insofar as genomic predictors are used for early diagnosis and prevention, such differences in prediction accuracy will lead to inequities in care. The loss in prediction accuracy across ancestry groups is believed to arise primarily from differences in patterns of linkage disequilibrium and allele frequencies at the common variants used in genome-wide association studies^16–18^. This realization has spurred a concerted effort to improve genotype imputation and develop trait prediction in a more diverse set of ancestries (e.g., in refs. ^19,20^).

An analogous argument about the importance of ancestry is often made when the goal is not to enable complex trait prediction, but instead to identify highly penetrant, severe disease variants. Here, the goal is often to identify genes whose disruption causes disease and ultimately learn about disease mechanisms, often by conducting burden tests or other association tests (see, e.g., Spence et al.^21^). In this case, the aim is to find as many *copies of pathogenic variants* in a gene as possible. To boost power, these pathogenic variants are often grouped into sets with similar functional effects: for example, loss of function alleles (LOF) of a given gene. Alternatively, researchers may want to establish an exhaustive *list of pathogenic variants*, for diagnostic purposes^22,23^. In these contexts, a commonly articulated concern about the disproportionate focus on people of recent European ancestry is that it will miss variants that are at appreciable frequency in other ancestries but rare in Europeans (e.g., ^11,22,24–27)^. Often provided as examples are genes that adapted in response to local environments, notably to geographically-restricted pathogens. In this view, there is a need for catalogs of “ancestry-specific” pathogenic variants, a term increasingly in use in the literature^28^.

Here, we argue that when the goal is to identify strongly deleterious alleles, the focus on ancestry groups is misguided. In point of fact, if we assume fitness effects of highly pathogenic mutations are shared across present-day environments, we should expect their frequencies to differ little among ancestries–much less so than variants of little or no fitness consequence (Charlesworth and Charlesworth^29^, Chapter 4). Indeed, although deleterious alleles are constantly being removed from the population by natural selection, they are repeatedly reintroduced by mutation, and should therefore persist at relatively stable, low frequencies^29^. As we show, the data conform to theoretical expectations: with very few exceptions, strongly deleterious mutations are similarly rare in all ancestry groupings and discovered at comparable rates in people of different ancestries. Thus, in the search for severe disease mutations, the emphasis on ancestry is unhelpful. In our view, it also comes at a potential cost, by conveying the erroneous impression that the causal mechanisms that relate mutations to disease outcomes differ across ancestry groups.

### Predictions

Much of the variation in the human genome is likely neutral or very close to neutral^30–32^. Accordingly, much of our intuition about genetic variation is based on models of genetic drift. Under such models, we expect alleles to vary in frequency across the globe by chance, and thus to differ across ancestry groupings. In some groups, an allele will have drifted up in frequency relative to others and so be more easily found by sequencing a relatively small sample. This effect is not large, as levels of genetic differentiation among human ancestry groups are relatively small^33–35^. Nonetheless, for a given number of genomes, more alleles will be found in more diverse sets of samples. These considerations informed early human population genomics projects such as the 1000 genomes project ^34^, which aimed to identify as many variants as possible by sequencing samples from many different ancestries and geographic locations.

When it comes to deleterious alleles, however, this intuition can break down. The frequency of a deleterious allele reflects a balance between the rate of mutations that give rise to it, genetic drift, and natural selection. The standard model for this “mutation-drift-selection balance” was developed almost a century ago^36–39^ and underlies recent efforts to identify constrained genes and sites from genomic variation data (e.g., in refs. ^7,40–46)^. The model predicts that if heterozygous carriers of an allele experience a fitness cost (i.e., if the mutation is “semi-dominant”), the population frequency of the allele will be close to *u*/(*hs)*, where *u* is the mutation rate to the disease allele and *hs* is the fitness effect of heterozygotes (reviewed in Fuller et al.^40^). For instance, if the mutation rate to any LOF allele in a gene is ∼10^-6^ per meiosis and the fitness cost of a LOF in heterozygotes is 1%, as is typical in humans^7,41–43^, then an allele that leads to the LOF of the gene should be found in 1 in 10,000 genomes on average (or roughly 1 in 5,000 people). Moreover, for very strong selection, the number of LOF alleles in a gene will be roughly Poisson distributed with mean *u*/*hs* in the population^42^; thus, in our example, given 100,000 people, the number of copies of a LOF mutation will usually lie between ∼4 and 16. Importantly, these predictions hold for all ancestries, regardless of their recent demographic history, so long as the fitness effects are the same^47,48^. Therefore, for semi-dominant mutations, our expectation should be that highly pathogenic alleles will vary little in frequency across ancestry groups.

This expectation will not necessarily hold if the disease mutation only has a fitness effect in homozygotes (or compound heterozygotes). In that case, the expected frequency of the allele may be higher, due to genetic drift, and greater differences may be found among ancestry groups, depending on their demographic histories^47,49^. Nonetheless, complete recessivity in fitness–as distinct from clinical presentation–is thought to be rare^12,50–52^. Another scenario in which we might expect greater differences among ancestry groups is under local adaptation^53^. The best-studied cases are alleles that were favored in response to the high endemicity of specific pathogens, but for which homozygosity leads to severe disease (e.g., those in refs. ^22,54–58)^. Strong positive selection at single loci is believed to be uncommon^59–61^, however, so these cases may be exceptions rather than illustrations of the more general ancestry distribution of pathogenic mutations^62^.

### Empirical findings

We tested these predictions using data from gnomAD release 4.1, a public resource that collates data from many different sources, and relied on their ancestry groupings, which are based on genetic similarity. Specifically, a person is assigned to a cluster by a classifier that is applied to the genetic principal components; the clusters are designated ancestry groups and are labeled on the basis of similarity to an existing reference panel of samples^63^. We consider the four clusters that include over 30,000 people other than “admixed American”, namely those labeled as having either recent non-Finnish European (NFE), South Asian (SAS), African or African-American (AFR), or Finnish ancestries (FIN). We note that although these labels are geographic, the clusters to which they refer sometimes regroup people from disparate locations: for example, AFR includes individuals identified as Puerto Rican and Qatari, as well as African^63,64^.

Focusing on synonymous variants, many of which are likely neutral, the average number of derived alleles per haploid genome is not significantly different across ancestry groups (Figure 1; see Supplementary Note 1 for details). This makes sense: assuming a fixed mutation rate, the number of neutral derived alleles carried by a genome reflects the time to the most recent common ancestor of humans, which is the same regardless of ancestry^65^. Similarly, the average numbers of missense and LOF alleles per genome are highly comparable across ancestries (Figure 1; Supplementary Note 1). These findings confirm what was found for smaller samples of thousands, and agree with theoretical expectations under mutation-selection-drift balance for semi-dominant mutations^47,49,66^. Genetic load, measured in this way, is remarkably similar regardless of ancestry^48,66^.

**Figure 1.**
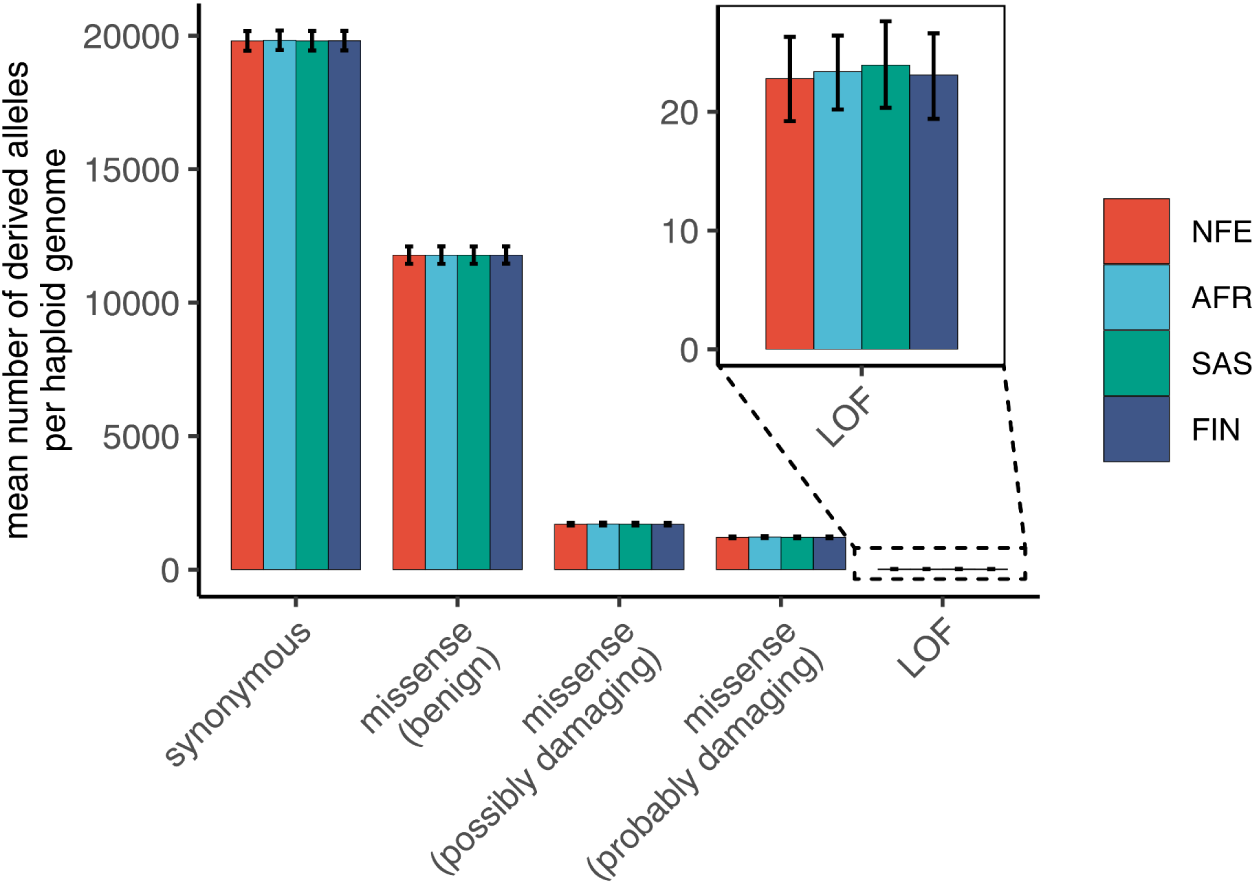
The average number of derived alleles per haploid genome does not differ between ancestry groupings. Abbreviations correspond to samples of Non-Finnish European (NFE), African or African-American (AFR), South Asian (SAS) and Finnish (FIN). Shown is the mean number of derived alleles of a given type (e.g., synonymous) per haploid genome (excluding derived alleles fixed in all four group samples). Only single nucleotide polymorphisms are included (i.e., no indels). The functional classification of variants is based on the VEP annotation in gnomAD v.4.1^63^. Only high confidence LOFs are shown, based on the LOFTEE annotation; the functional classification of missense variants is based on PolyPhen-2^67^. All groups are downsampled to 60,000 chromosomes, by binomial sampling at each site (see Supplementary Note 1). Variants are polarized based on the ancestral sequence inferred in 1000 Genomes Project^34^ and the functional annotation is corrected for reference bias following Simons et al.^47^. Given that individual data are not available for gnomAD, confidence intervals are estimated using bootstrap resampling of genes (see Supplementary Note 1).

Although the mean number of derived alleles per genome is the same, the frequency distribution of alleles depends on the demographic history of each group^47^. For example, due to the Out-of-Africa bottleneck, samples of non-African ancestry have a higher proportion of variants segregating at appreciable frequencies and a deficit of the rare alleles found in sub-Saharan ancestries^47,68,69^. Therefore, if an allele frequency cutoff is imposed, for instance to decrease the chance of mis-annotating a LOF allele, it will affect distinct ancestry groups differentially and may lead to apparent differences in genetic load.

LOF alleles are a relatively well defined set of pathogenic alleles on which to train our intuition. Of the different functional annotations considered, they are most likely to lead to lethality or severe developmental disorders^43,70–73^, and are subject to purifying selection that is strong relative to drift: on average, a mutation that leads to the loss of a gene copy is estimated to inflict an average fitness cost of 1% ^7,41–43^. To hone in on the subset that are likely to be most deleterious, we focus on strongly constrained genes for which the loss of a single copy is predicted to inflict a fitness cost of >5%, based on features such as gene expression patterns across tissues, protein structure and evolutionary conservation, but importantly not on allele frequency information (i.e., using the prior means reported in Zeng et al.^43^). In such genes, the number of LOF alleles differs very little between pairs of ancestry groups (matched for sample size), almost always by fewer than 10 copies out of 60,000 chromosomes (Fig. 2). This holds true whether the label regroups a set of more heterogeneous ancestries (e.g., AFR) or a more isolated, “founder population” (FIN)一that is regardless of recent demographic history. This observation indicates that alleles that are highly deleterious in people in one ancestry grouping tend to be so across the distributions of genetic backgrounds and environments of other ancestries as well.

**Figure 2.**
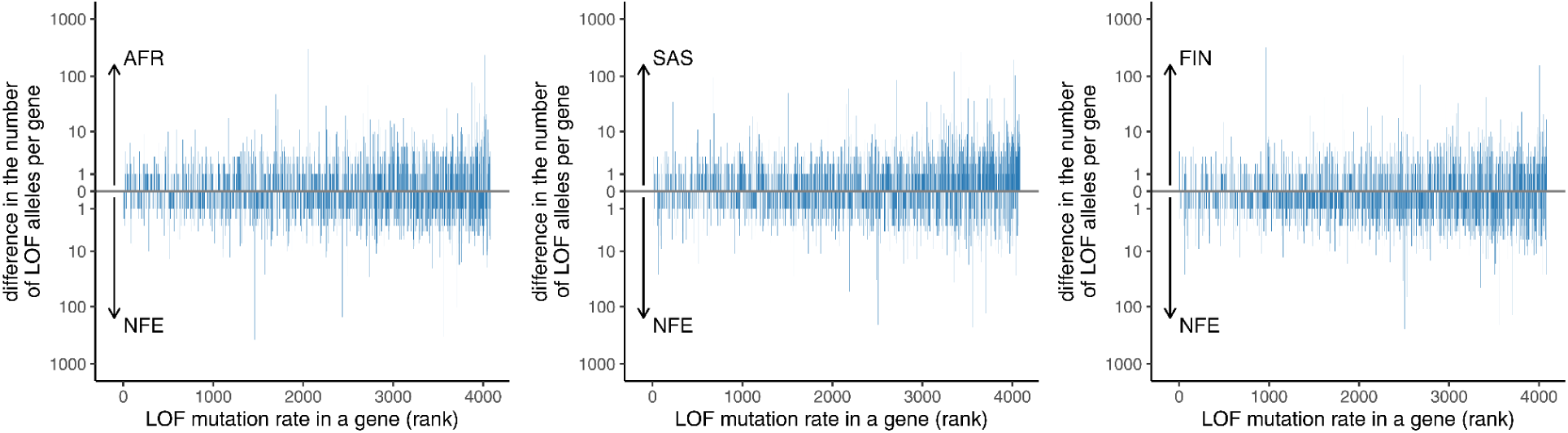
The number of LOF copies differs little between ancestry groups in strongly constrained genes. We consider strongly constrained genes for which the mean prior *hs* is >5% ^43^. Genes are sorted by their expected LOF mutation rate^45^. Both groups are downsampled to 60,000 chromosomes, by binomial sampling at each site (see Supplementary Note 1). The allele counts of all derived LOF variants in a gene are summed to obtain the total number of LOF alleles in a gene. LOFs with derived allele frequency above 1% were excluded to avoid misannotation; this cutoff led to the removal of four LOF variants, and of these only one varied markedly in frequency between groups. In the absence of an allele frequency cut-off, the total number of highly deleterious LOF alleles does not differ significantly between groups (bootstrapping genes, p ≥ 0.24; see Supplementary Note 1 for details).

Differences between ancestry groups in the number of highly deleterious LOF alleles per gene are well predicted by the mutation-drift-selection balance model: comparing NFE and AFR for example, only 4% of genes (154 out of 4072) fall outside the 95% CI (Figure 3A), and the average absolute value of the difference in allele numbers between groups is close to its expectation (observed/expected = 0.91; 0.95 without an allele frequency cut-off). Overall, highly deleterious LOF alleles are rare in all ancestries, and tend to differ little in frequency among groups on the gene level.

**Figure 3.**
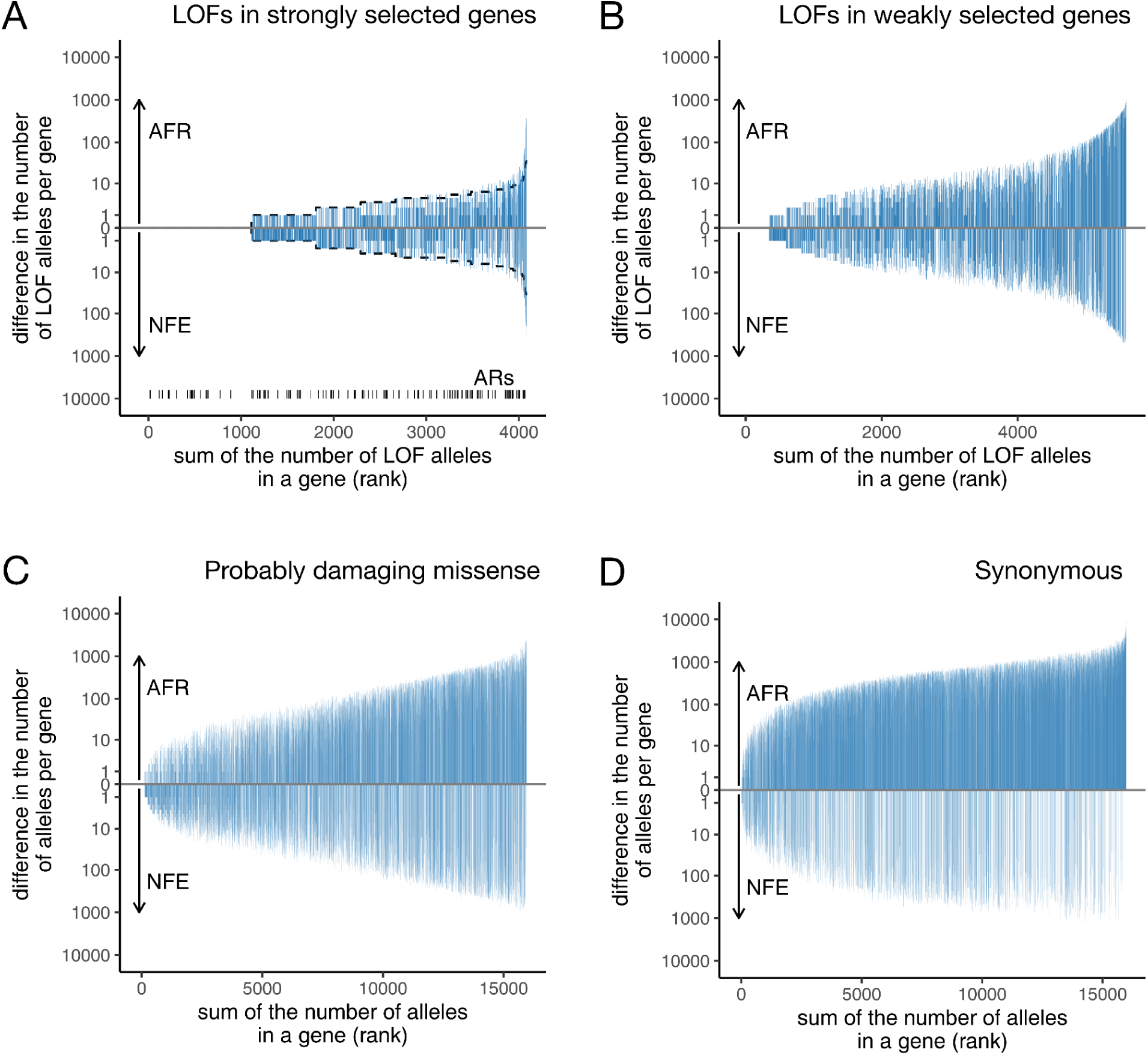
Strongly deleterious alleles show the smallest difference between the number of derived alleles in AFR and NFE groupings. Genes are sorted by the total number of derived alleles observed in the gene in the joint sample of AFR and NFE. AFR and NFE are downsampled to 60,000 chromosomes. The allele counts of all derived variants of a given class in a gene are summed to obtain the total number of alleles in a gene. Variants with derived allele frequency above 1% were excluded to avoid misannotation of LOF variants, and for comparability to missense and synonymous variants; this cut-off induces an asymmetry between ancestry groups for nearly neutral and weakly deleterious alleles (see Supplementary Note 2). **(A)** High confidence LOFs in strongly selected genes, i.e., genes with mean prior estimate of hs > 5% for LOFs^43^. The dashed lines show the predicted 95% interval for the difference between the number of copies in AFR and NFE, assuming that they come from the same Poisson distribution (with a mean given by the sum of the copies in each population). These predicted intervals are only shown for panel A, as they are based on a model of mutation-selection balance for strongly deleterious alleles. The autosomal recessive genes are shown with tick marks at the bottom, and are classified based on an annotation of OMIM disease genes^74,75^. **(B)** High confidence LOF variants in weakly selected genes, i.e., genes with mean prior hs < 0.05% for LOFs^43^. **(C-D)** Probably damaging missense and synonymous variants in all genes. The functional impact of missense variants is from PolyPhen-2^67^. The annotations of derived LOFs and missense variants were corrected for reference bias following Simons et al.^47^; for additional details, see Supplementary Notes 2 and 3. Plots for SAS and FIN populations versus NFE are in Supplementary Figures 1 and 2; plots without filtering for variant frequencies are in Supplementary Figures 3-5.

The subset of strongly selected genes classified by clinicians as autosomal recessive tend to have higher LOF frequencies in a combined sample of AFR and NFE (Wilcoxon rank-sum test p-value < 1e-5; Figure 3A and Supplementary Figures 1A and 2A for other ancestry groups), as expected from mutation-selection-drift balance for recessive alleles. Moreover, the number of LOF allele copies differs more between ancestry groups for this subset of 117 genes than for other strongly selected genes^47,49^. Nonetheless, differences in the number of copies remain small (Figure 3A).

Again as expected from theory, the more deleterious the fitness effect, the smaller the difference in the frequency across ancestry groups: thus, LOF mutations that are predicted to strongly deleterious differ least in frequency among ancestry groups, more weakly deleterious LOF more so (Figure 3B), and probably damaging missense and synonymous mutations yet more (Figure 3C-D, Supplementary Figures 1 and 2). These findings imply that contrary to what a few well-studied examples of more weakly deleterious variants suggest, we should not expect highly deleterious mutations to be absent or ultra-rare in European samples but common in non-European ancestries. To evaluate this claim explicitly, we consider how commonly a gene has no LOF variants in a 60K chromosome sample of NFE ancestry, but carries LOFs at appreciable frequency in the same sample size of other ancestries. It is exceedingly uncommon: none of the 1634 genes with no LOFs observed in the NFE subsample have a LOF frequency above 0.1% in any of the ancestries considered (Supplementary Figure 6). A similar pattern is observed at the level of individual LOF variants (Supplementary Figure 7). For example, in AFR, among all strongly selected LOF variants absent in the NFE sample but present in the AFR one, only 0.1% have a frequency above 0.1%.

In contrast to strongly deleterious LOF mutations, missense alleles classified as “probably damaging” are more likely to be absent in one sample but at non-negligible frequencies in other groups: as an example, 2% of probably damaging missense mutations that are not found in the NFE sample occur in AFR at frequencies above 0.1% (Supplementary Figure 7). Such missense alleles usually have a smaller effect on disease risk than do gene losses and are also less likely to be strongly deleterious^7,21,76^. As a result, they may reach appreciable frequency in an ancestry group and contribute non-negligibly to the variance in disease risk^77^. However, when an allele is frequent in an ancestry group, it will typically be widely shared across ancestries^78^.

Because the frequencies of weakly selected alleles are more subject to genetic drift than are strongly selected alleles, power may be gained by performing association tests in groups in which a disease allele has by chance increased in frequency, and in which phenotypic effects are more easily assessed, such as in founder populations (e.g., in refs. ^79,80^). Recently bottlenecked populations have an increased chance of revealing at least one trait-associated variant that has drifted up to higher frequency (e.g., Lynch et al.^81^). But the mean frequency of pathogenic alleles is expected to be no higher there than in other groups (see, e.g., FIN in Figure 1; refs. ^47,66^), as other disease-associated alleles will have drifted down in frequency. Therefore, while mapping studies in ancestry groups that have experienced substantial genetic drift may be helpful in identifying weakly deleterious, semi-dominant alleles or strongly deleterious, recessive alleles associated with disease risk, in the absence of prior evidence of higher prevalence of a particular disease due to genetic effects (rather from environmental exposures), the optimal study design needs to be evaluated explicitly.

Next, we assess how many additional copies or loci could be identified by sequencing individuals of different ancestries, compared with sequencing more individuals of the same ancestry. To this end, we assume that 60K haploid genomes of recent NFE ancestry have already been sequenced, and ask how many new variants would be discovered by sequencing an additional 60K chromosomes from either the same or distinct ancestry groups. For all types of mutations, sequencing other ancestry groups leads to the discovery of a similar number of *copies* as increasing the sample size of NFE (Figure 4A; see Supplementary Figure 8A-C for analogous results for other ancestries, as well as Steiner et al.^48^ Figure 6), consistent with the deleterious mutation load per genome being the same in all ancestry groups^47^.

**Figure 4.**
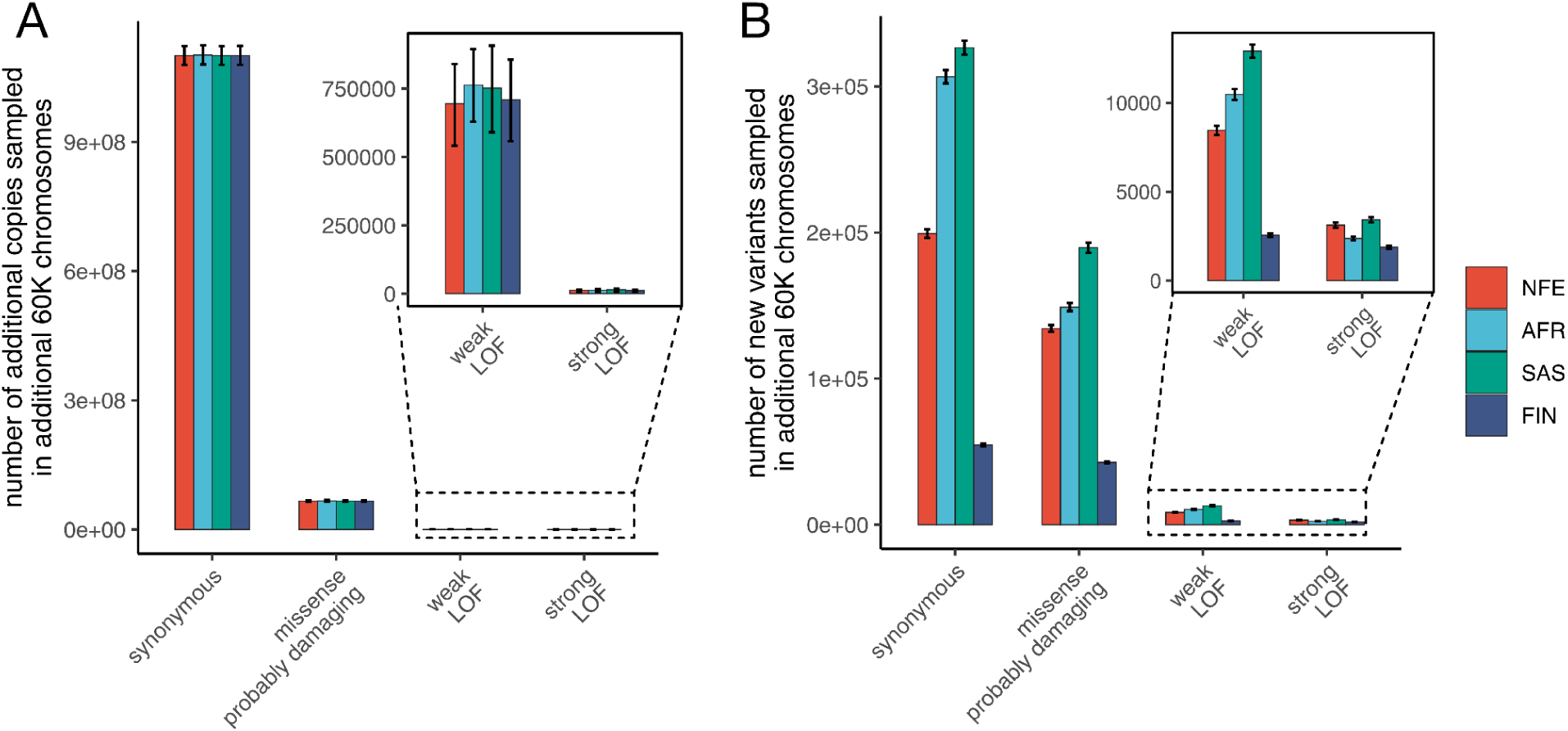
Sampling additional samples of any ancestry yields similar numbers of strongly deleterious variants. **(A)** The number of additional copies of derived alleles of a given functional class sampled by sequencing an additional 60,000 chromosomes from a given ancestry group in addition to 60,000 chromosomes from NFE. **(B)** The number of new variants sampled by sequencing an additional 60,000 chromosomes. Annotations for probably damaging missense mutations are based on PolyPhen-2^67^, high confidence LOF variants with prior hs < 0.5% are referred as weak and with hs > 5% as strong^43,45^. The 95% confidence intervals are estimated from 10,000 bootstrap resamples of genes (see Supplementary Note 1).

If we instead ask how many *newly discovered variants* are found by sequencing different ancestries, the findings are not identical across groups (Figure 4B, Supplementary Figure 8D-F, as well as Steiner et al.^48^ Figure 6). Considering synonymous variants, which are likely closest to neutrally evolving, sequencing AFR and SAS populations yields the most new variants, whereas sequencing a genetically more similar and less diverse FIN cohort results in fewer new variants. Viewed in terms of the underlying ancestral recombination graph in which all the samples are embedded, adding individuals with less similar ancestries adds somewhat longer lineages on average and thus is more likely to reveal additional mutations^65,82^. One implication is that, as expected from theory, sampling sets of individuals with diverse ancestries is a more efficient way of building up a repository of neutral variation^34,61,63,83^.

The same is not true for highly pathogenic mutations, however. Variant discovery rates differ little by ancestry group for LOFs in genes under strong selection: an illustration, while sequencing an additional 60K NFE samples leads to the identification of 3005 new variants, on average, 2525 new variants are found by instead sequencing one of the other three ancestry groups (see also Steiner et al.^48^, Figure 4A). This again makes sense: highly pathogenic variants are recent, meaning that they are found towards the tips of the lowest branches of the ancestral recombination graph; such tips are added by sampling people of any ancestry.

For a related reason, the rate of discovery of highly pathogenic variants is likely to be strongly influenced by study design. For example, when datasets include extended family members, the same pathogenic variant is more likely to be observed multiple times (for an analogous spatial argument, see ^48^). The extent to which different recruitment strategies influence discovery rates within and between ancestry groups remains to be evaluated.

It is also important to note that even when a variant has of yet only been discovered in one ancestry grouping, it is not “ancestry-specific.” Even at the level of a single base (e.g., a specific non-synonymous mutation), a given mutation will have arisen hundreds of times independently in the last few generations, given >8 billion humans at present and a mutation rate per base pair of ∼10^-8^ per generation. In the population of India alone, a rough calculation suggests that a single nucleotide allele will have arisen ∼10 times in the last generation on average. Therefore, a given mutation will be found in any sufficiently large set of humans. With strong selection against it, its frequency will be similar across ancestry groups. Indel variants defined by their exact breakpoints arise at a lower rate than point mutations^84^ and will thus show lower recurrence across groups. However, as a class, indels in a gene with severe pathogenic consequences are expected to be at comparable frequencies across ancestry groups, as is the case for LOF variants.

We note further that because highly deleterious alleles will tend to be young, they will be geographically-restricted^48^. For example, a rare pathogenic allele found in the UK Biobank on a particular haplotype may only be shared among people in the North-East of England; meanwhile, the pathogenic allele is expected at similar frequencies across Europe–indeed the world–on many different haplotypes. The recency of highly pathogenic mutations carries another implication: the designation of a disease mutation found in a United States patient as, e.g., AFR based on the person’s ancestry may be inaccurate. A variant that inflicts a fitness effect of, say, 5% in heterozygotes is typically only five generations old (Supplementary Note 4) and will be found identical by descent only in close relatives of the carrier. The description of the allele as “African-specific” would therefore not be informative about where it arose, or where more carriers might be found. Therefore, even a pathogenic mutation that is shared identical by descent is not the property of a major ancestry group; rather, it is an allele found in an extended family.

In humans, as in other species, population structure exists at every scale of resolution. Ancestry groupings are an approximation at a broad scale, which can be useful for some analyses^85,86^. Notably, in the study of common variants and their roles in complex disease susceptibility, it can be helpful to cluster individuals and combine results across individuals who are more genetically similar to each other (in terms of their allele sharing and patterns of linkage disequilibrium), motivating the use of such ancestry groups. In searching for rare, deleterious variants that are young and geographically-restricted, by contrast, the emphasis on ancestry groups in study design and interpretation is unfounded.

## Conclusion

Given that highly pathogenic mutations recur frequently by mutation and are at similar low frequencies in all ancestry groups, rates of discovery will be similar regardless of ancestry. These considerations clarify that disease variants are to be found in extended families, not genetic ancestry groupings, and the information that they provide about human biology is not specific to an ancestry. Genes with highly pathogenic mutations in one ancestry group illuminate causal pathways underlying disease outcomes in members of all groups: regardless of the ancestries sampled, once enough copies of a pathogenic allele have been found to establish a link to disease risk, it is clear that the gene is of functional relevance for pathophysiology. Similarly, we do not need to sequence millions of people from every ancestry group to learn that a mutation that has never been observed in huge samples is more likely to be deleterious, or inversely, that a mutation at high frequency is unlikely to be highly pathogenic.

There are many important reasons to diversify study participants–not to mention researchers–in medical research. In human genetics, mapping studies may gain power from the distinct distributions of environmental exposures experienced by people from different ancestry groups or in the case of more weakly deleterious alleles, by leveraging their somewhat distinct demographic histories. When the goal is to identify strongly deleterious mutations, however, the focus on genetic differences among ancestry groups is misguided, and obscures the fact that the causal mechanisms underlying disease risk are shared among people of all ancestries.

## Acknowledgments

We thank Ipsita Agarwal, Michael “Doc” Edge, John Novembre, and Jonathan Pritchard for helpful comments on a draft of the manuscript.

## Supplementary Notes

### 1. Assessments of statistical significance

As a measure of genetic load, we focused on the average number of derived alleles per haploid genome (see Simons and Sella^66^ and references therein, for a discussion of the various choices of load measures). Given that gnomAD does not provide haplotypes, to obtain this average, we estimated the frequency of each allele in 60,000 chromosomes, by sampling from a binomial distribution with mean the derived allele frequency within that ancestry group. Each variant was estimated independently, assuming no linkage disequilibrium (LD) between them. To obtain the average allele count per haploid genome, we summed the number of alleles of a given functional class across the group and divided by 60,000. We excluded genes in which LOF variants are predicted to drive clonal expansions in spermatogonia, increasing their likelihood of transmission to offspring^87^. In total, 14 such genes were removed from the analysis: 12 genes were excluded among those with a prior mean *hs* above 5% (Figures 2 and 3A) and none from genes with a prior mean *hs* below 0.5% (Figure 3B)^43^.

To estimate confidence intervals for the mean number of derived alleles per haploid genome (Figure 1), we bootstrapped genes (the same procedure was used for Figure 4). To that end, we generated 10,000 resampled datasets, each the same number of genes as the original, by randomly selecting genes with replacement and calculating the total number of derived alleles for each functional class (i.e., high-confidence LOFs, missense variants annotated as probably damaging, possibly damaging, or benign, and synonymous variants). From these resamples, we estimated the 95% quantile intervals shown in Figure 1. We also used the bootstrap samples to compare the number of alleles per genome in the NFE to other groups; comparisons are not statistically significant after correction for multiple tests (p>0.04; see Supplementary Table 1).

We note that bootstrapping individuals rather than genes (as done in Simons et al.^47,49^ and Do et al.^47,49^) may be a more appropriate approach for testing significant differences in genetic load between ancestries, because it better mimics the randomness of drawing samples from a large number of people assigned an ancestry label. However, that is not feasible here, as gnomAD only provides allele frequencies, not individual genotypes. Moreover, a limitation of bootstrapping individuals is that they share genealogical histories and are not independent, all the more so in large sample sizes ^47^. Instead, bootstrapping genomic regions (here, genes) may better capture uncertainty due to evolutionary histories (assuming no linkage disequilibrium between variants in different genes, as seems plausible). Regardless of the exact method used to assess the confidence intervals, the counts of alleles are qualitatively across ancestries (as can be seen in Figure 1 and Figure 4A).

### 2. Effect of allele-frequency cutoff

In analyses presented in the main text, we excluded variants with derived allele frequency above 1% to avoid mis-annotating LOF variants^43,71^; for comparability, we then imposed the same cut-off on missense and synonymous variants. The differences between the number of derived alleles between groupings without an allele frequency threshold are shown in Supplementary Figures 3-5.

Differences in demographic histories, e.g., due to the Out-of-Africa bottleneck, lead to distinct allele frequency distributions across ancestry groups, and hence distinct effects of the cut-off. For instance, in the Finnish ancestry group, which is often described as a founder population ^88^, there is an enrichment of higher frequency variants compared to NFE and the allele frequency cutoff thus has a larger effect on the number of carried derived alleles (Supplementary Figure 2). Similarly, the cut-off leads to the exclusion of more variants from the NFE than AFR (Figure 3B-D). Given that selection maintains strongly selected variants at similar frequencies in different ancestry groups, the cut-off has much less effect on LOF variants: it leads to the removal of only four variants, of which one varied markedly in frequency between groups. As expected, the effect of the cut-off is more pronounced for weakly deleterious and neutral variants.

### 3. Effect of discrete ancestry assignments

Without an allele frequency cut-off, there is a slight skew towards alleles being at higher frequency in non-NFE as compared to NFE in genes with relatively high LOF frequencies (Figure 3A and B, Supplementary Figures 1-5A and B). This asymmetry is more readily visualized in a scatterplot of the derived allele frequencies of LOF alleles in NFE versus AFR (Supplementary Figure 9A). There are vertical bands of points corresponding to alleles that are polymorphic in AFR and absent or at very low frequency in NFE becomes apparent, when there are no equivalent horizontal bands of alleles that are (near) absent in AFR but polymorphic in NFE. A clear diagonal band away from the 1-to-1 line is also present.

The cause of this asymmetry is likely to be the assignment of samples in gnomAD to discrete ancestry groupings. Assignment to a single ancestry, using a dataset with a high proportion of people of recent European ancestry, leads people with some NFE ancestry to be included in groups other than NFE, for example when African-Americans with recent European ancestry are included in the AFR group. As a consequence, rare alleles that are polymorphic in NFE end up polymorphic in the AFR grouping, even if they are extremely rare in people living in continental Africa. In contrast, very few individuals of non-European genetic ancestry are included in the NFE cluster, so the reverse very rarely occurs. Thus, the diagonal ridge parallel to the x=y line is populated by SNPs that are (near) absent in continental African samples in the 1KGP but present in the AFR sample (orange dots). This diagonal ridge corresponds to rare alleles in AFR that are at ∼15% of the NFE frequency, likely reflecting recent European ancestry in African-Americans that are included in the gnomAD AFR samples^89^. For most (75%) of AFR samples, no labels are provided; among existing labels are a disparate set of geographic locations, nationalities, races and ethnicities, including, e.g., African-American, African-Caribbean, Mexican, Puerto Rican, Qatari, White and Black African^63,64^. This effect of discrete ancestry assignments is not specific to recent admixture in the history of African-Americans; similar albeit weaker patterns are also seen in Supplementary Figures 9B-C, for example.

### 4. Our estimate of the age of segregating strongly selected, deleterious LOFs

To model LOF mutations in a gene, we used a forward, population-genetic simulation framework originally described in^47^ and modified in^41^. In this model, the gene evolves as a single non-recombining, biallelic locus under a mutation rate *μ* in a Wright-Fisher diploid population with an effective population size *Ne*; *μ* here is the mutation to a loss of function. The fitness of the genotypes are given by 1, 1-*hs*, 1-*s*. At most one mutation can arise each generation. We used the Schiffels–Durbin demographic model for population growth in Europe^90^, modified to include a larger recent effective population size ^41^. To determine the mean allele age, in each case in which the locus is segregating in the population at present, we recorded the generation in which the mutation arose. LOF allele ages were obtained from 10,000 forward simulations, using *μ* = 10^-6^ per locus per generation^41^, *h* = 0.5, and *s* = 10%.

## Supplementary Tables

**Supplementary Table 1.** The relative difference in the number of derived alleles for each ancestry group compared to NFE. All ancestry groups are downsampled to 60,000 chromosomes by binomial sampling at each site. P-values are calculated using bootstrap resampling of genes and adjusted for multiple testing by a Bonferroni correction (for 15 tests). For additional details, see Supplementary Note 1.

|  | AFR vs NFE | SAS vs NFE | FIN vs NFE |
| --- | --- | --- | --- |
| **synonymous** | +0.11% (p = 1.0) | +0.0% (p = 1.0) | +0.05% (p = 0.23) |
| **missense (benign)** | +0.04% (p = 1.0) | -0.01% (p = 1.0) | +0.02% (p = 1.0) |
| **missense (possibly damaging)** | +0.37% (p = 1.0) | +0.15% (p = 1.0) | +0.01% (p = 1.0) |
| **missense (probably damaging)** | +0.89% (p = 0.11) | +0.17% (p = 1.0) | +0.11% (p = 1.0) |
| **LOF** | +4.69% (p = 1.0) | + 6.64% (p = 0.042) | + 0.77% (p = 1.0) |

## Supplementary Figures

**Supplementary Figure 1.**
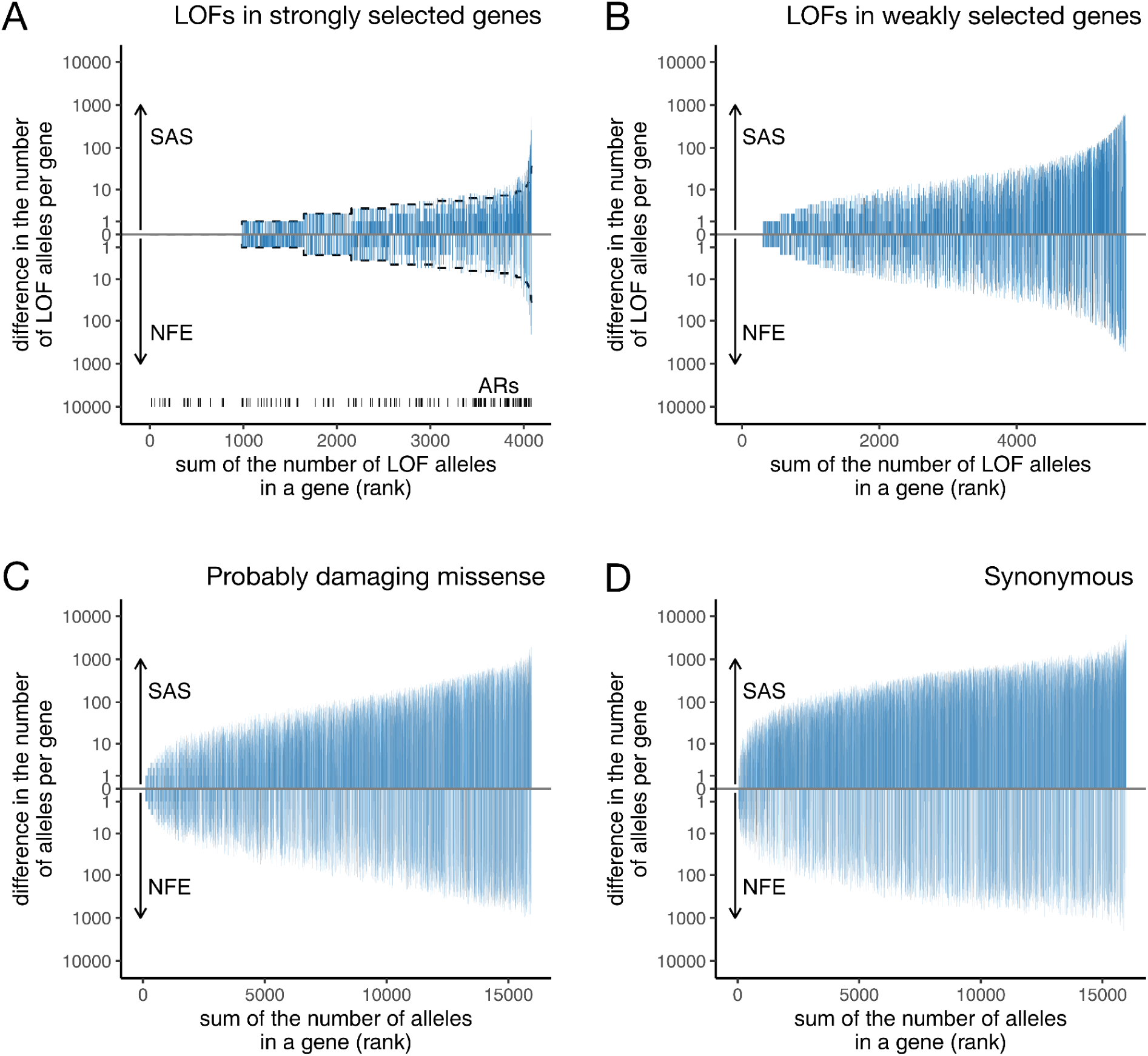
Difference between the number of derived alleles in SAS and NFE groupings, given a 1% allele frequency cutoff. Both groups are downsampled to 60,000 chromosomes. Genes are sorted by the total number of alleles observed in the gene in the joint sample. For further details, refer to the caption of Figure 3.

**Supplementary Figure 2.**
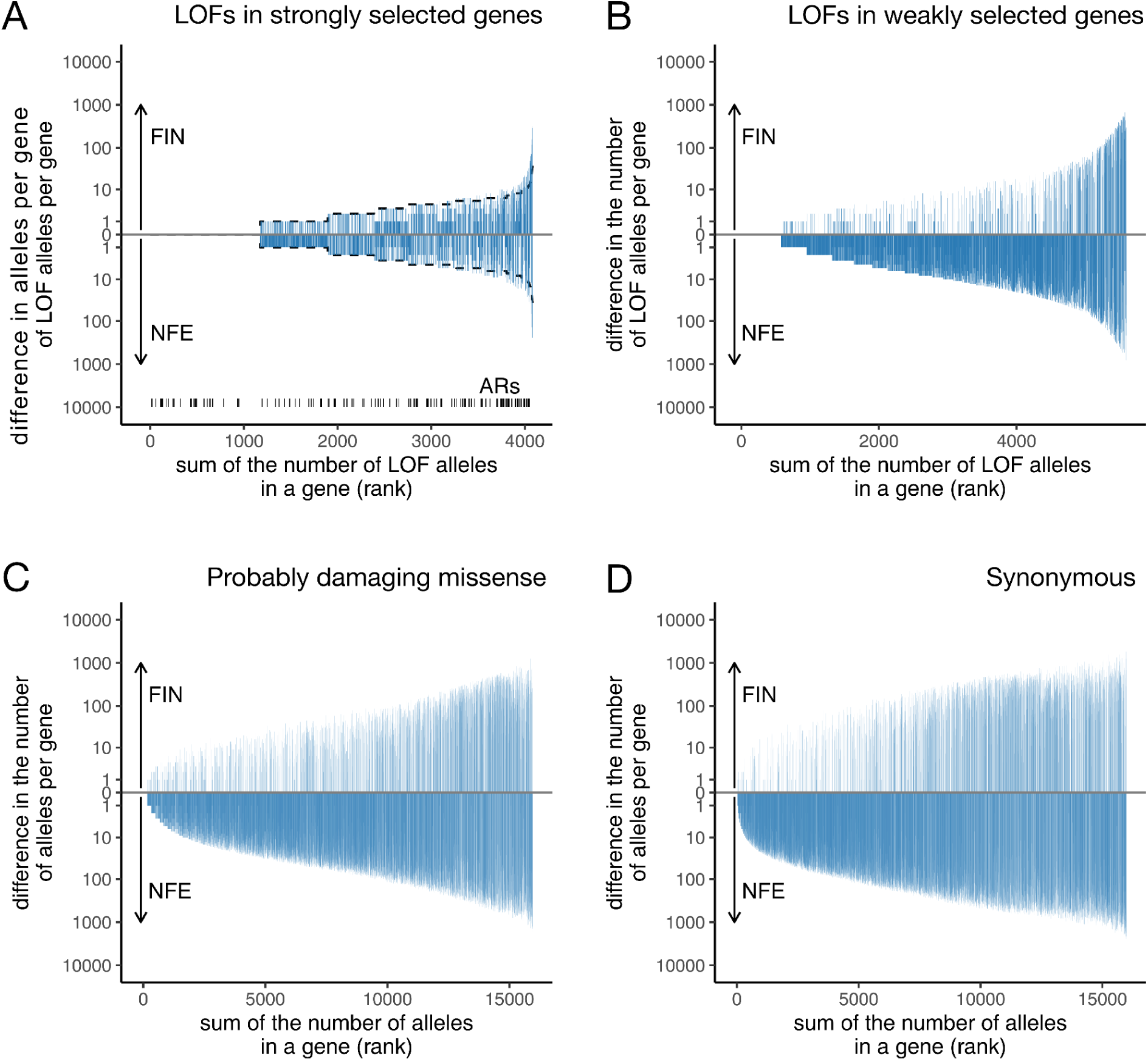
Difference between the number of derived alleles in FIN and NFE groupings, given a 1% allele frequency cutoff. Both groups are downsampled to 60,000 chromosomes. Genes are sorted by the total number of alleles observed in the gene in the joint sample. For further details, refer to the caption of Figure 3.

**Supplementary Figure 3.**
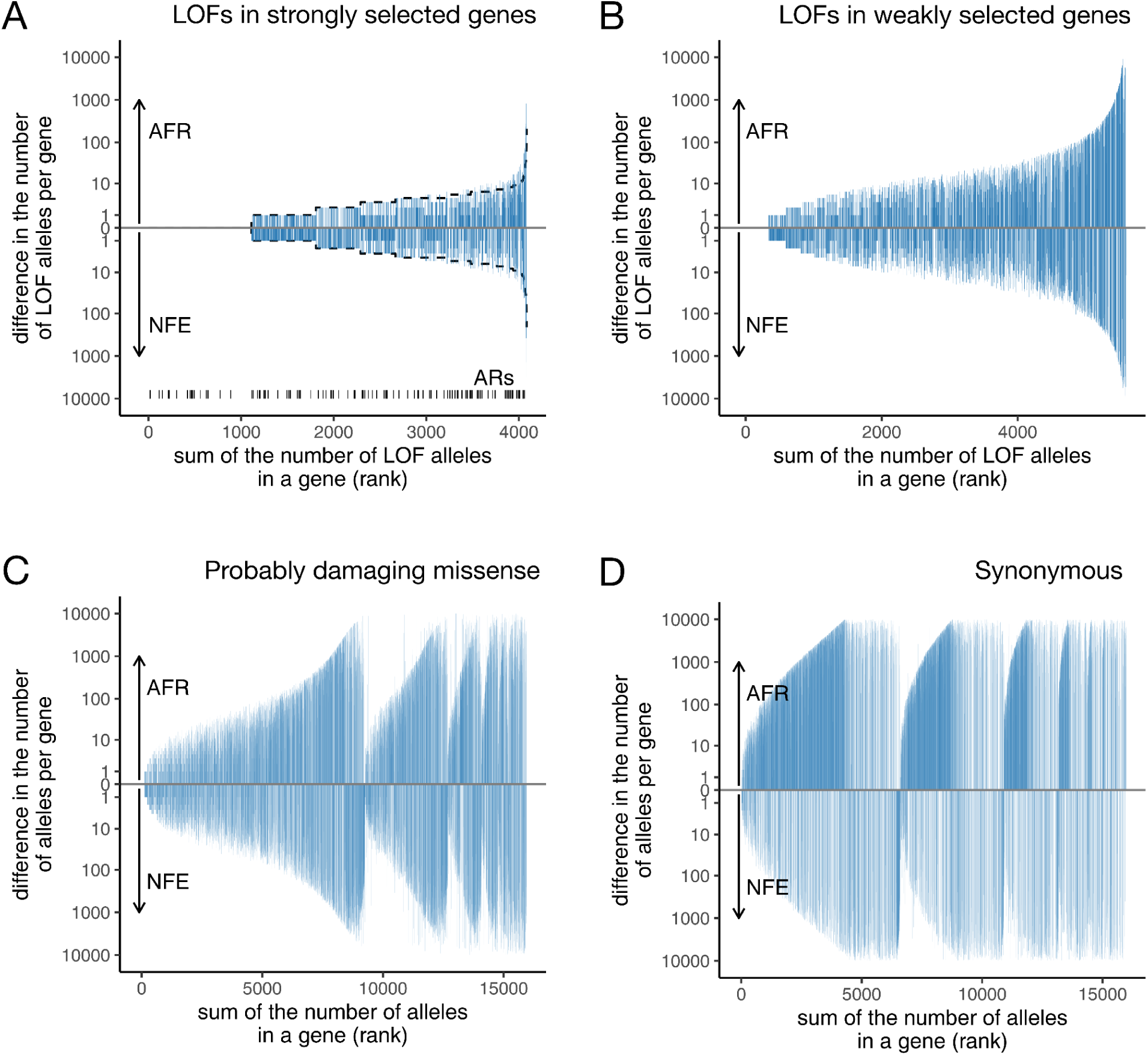
Difference between the number of derived alleles in AFR and NFE groupings *without* an allele frequency cutoff. Both groups are downsampled to 60,000 chromosomes. Genes are sorted by the total number of alleles observed in the gene in the joint sample. For further details, refer to the caption of Figure 3. The repeated drops in the difference between allele counts in (C) and (D) are due to variants at high but similar frequencies in both populations (i.e., those near fixation). The repeated drops in the difference between allele counts in (C) and (D) are due to variants that are shared between populations and have high but similar frequencies in both. These variants are likely identical by descent and are near fixation, meaning that the absolute difference in allele frequency between AFR and NFE is small even though the allele itself is present at a high frequency in both groups. Since genes are sorted by their total allele count, such genes are pushed to the right on the x-axis but remain close to zero on the y-axis, creating a non-monotonic pattern.

**Supplementary Figure 4.**
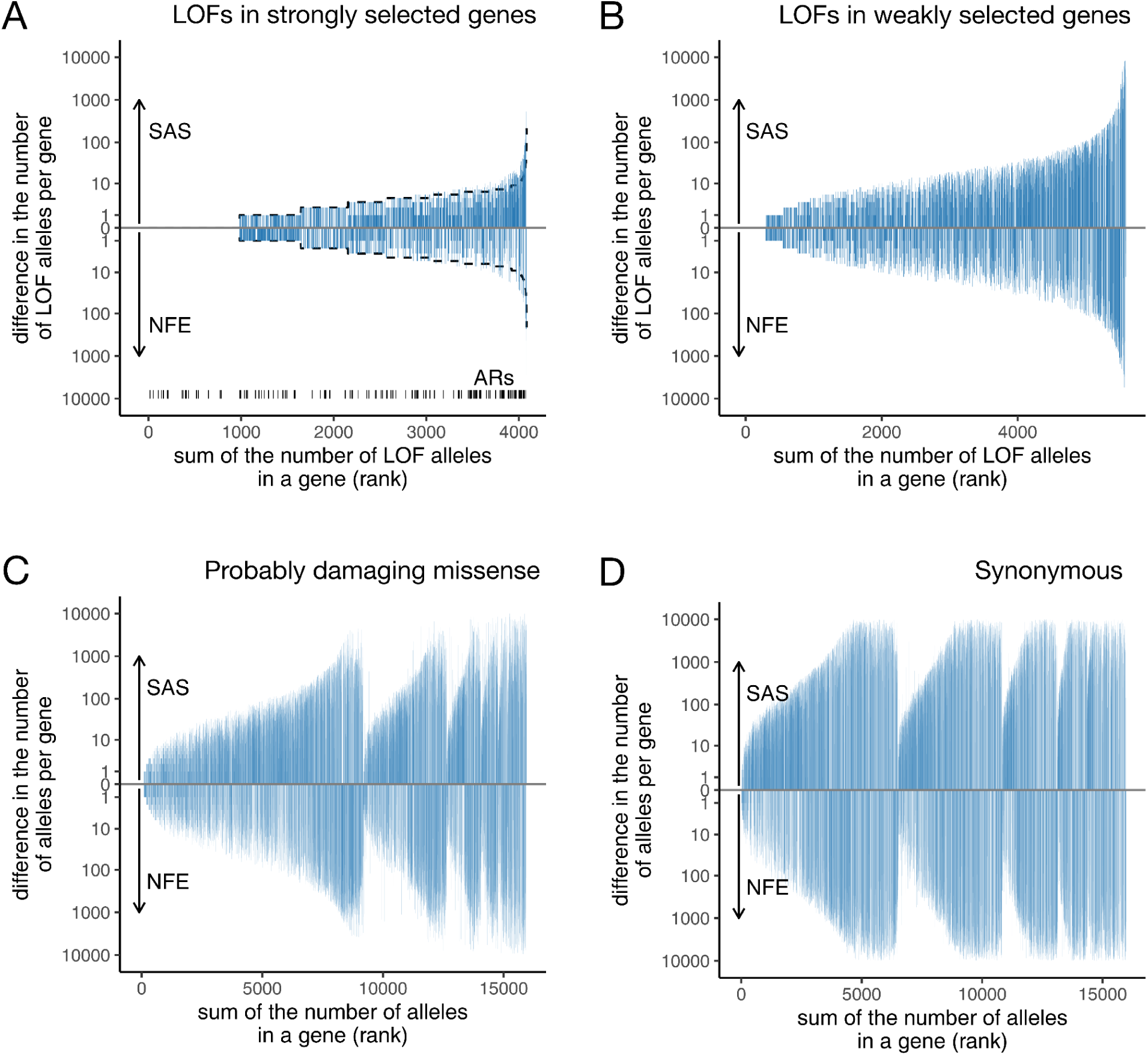
Difference between the number of derived alleles in SAS and NFE groupings *without* an allele frequency cutoff. Both groups are downsampled to 60,000 chromosomes. Genes are sorted by the total number of alleles observed in the gene in the joint sample. For further details, refer to the caption of Figure 3.

**Supplementary Figure 5.**
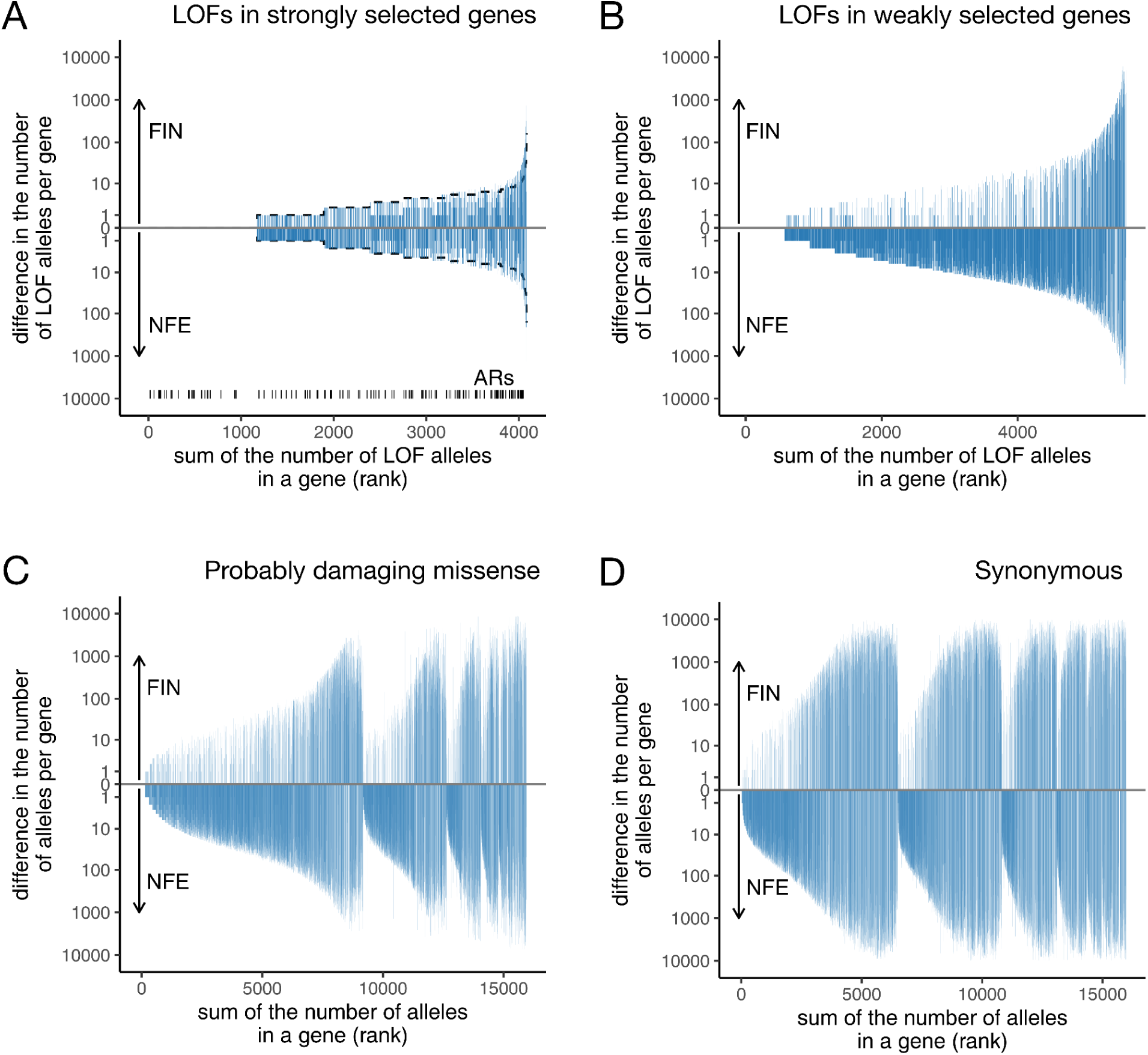
Difference between the number of derived alleles in FIN and NFE groupings *without* an allele frequency cutoff. Both groups are downsampled to 60,000 chromosomes. Genes are sorted by the total number of alleles observed in the gene in the joint sample. For further details, refer to the caption of Figure 3.

**Supplementary Figure 6.**
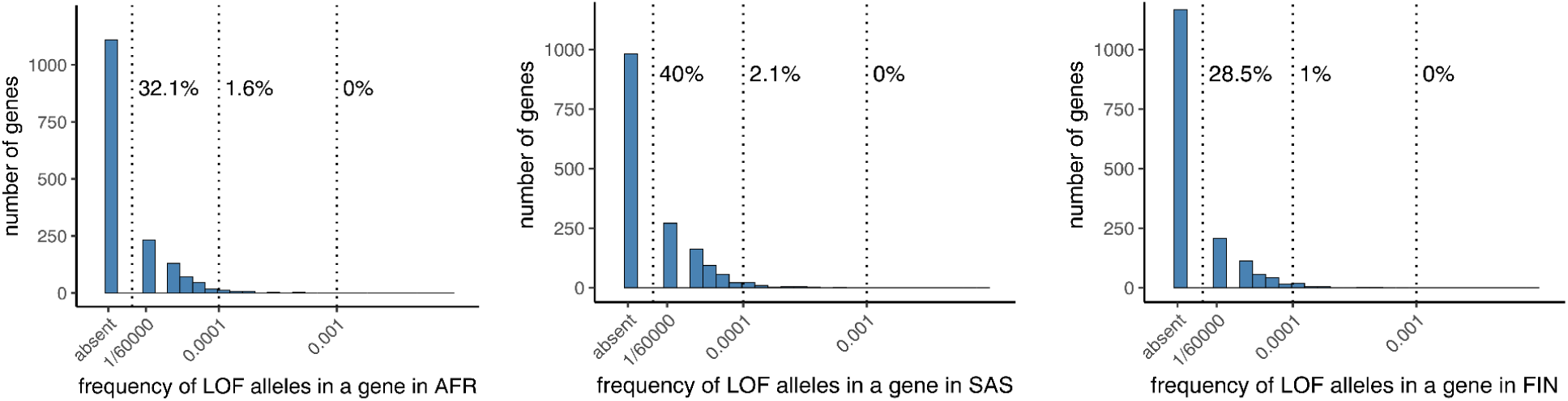
The number of strongly selected genes with LOF alleles observed in AFR, SAS and FIN ancestry groups, given that no LOF allele is found in NFE. All sample sizes were downsampled to 30,000 individuals (60,000 chromosomes). We focused on the 1634 strongly selected genes (i.e., with a mean prior hs > 5% from Zeng et al.^43^), for which no LOF is observed in the NFE cohort. The frequencies of all derived LOF variants in a gene are summed to obtain the total LOF frequency in a gene. The percentages show the fraction of genes with more than one copy of a LOF allele (>1/60,000), with LOF alleles at a frequency >0.01% in the focal ancestry group, and with LOF alleles at frequency >0.1% in the focal ancestry.

**Supplementary Figure 7.**
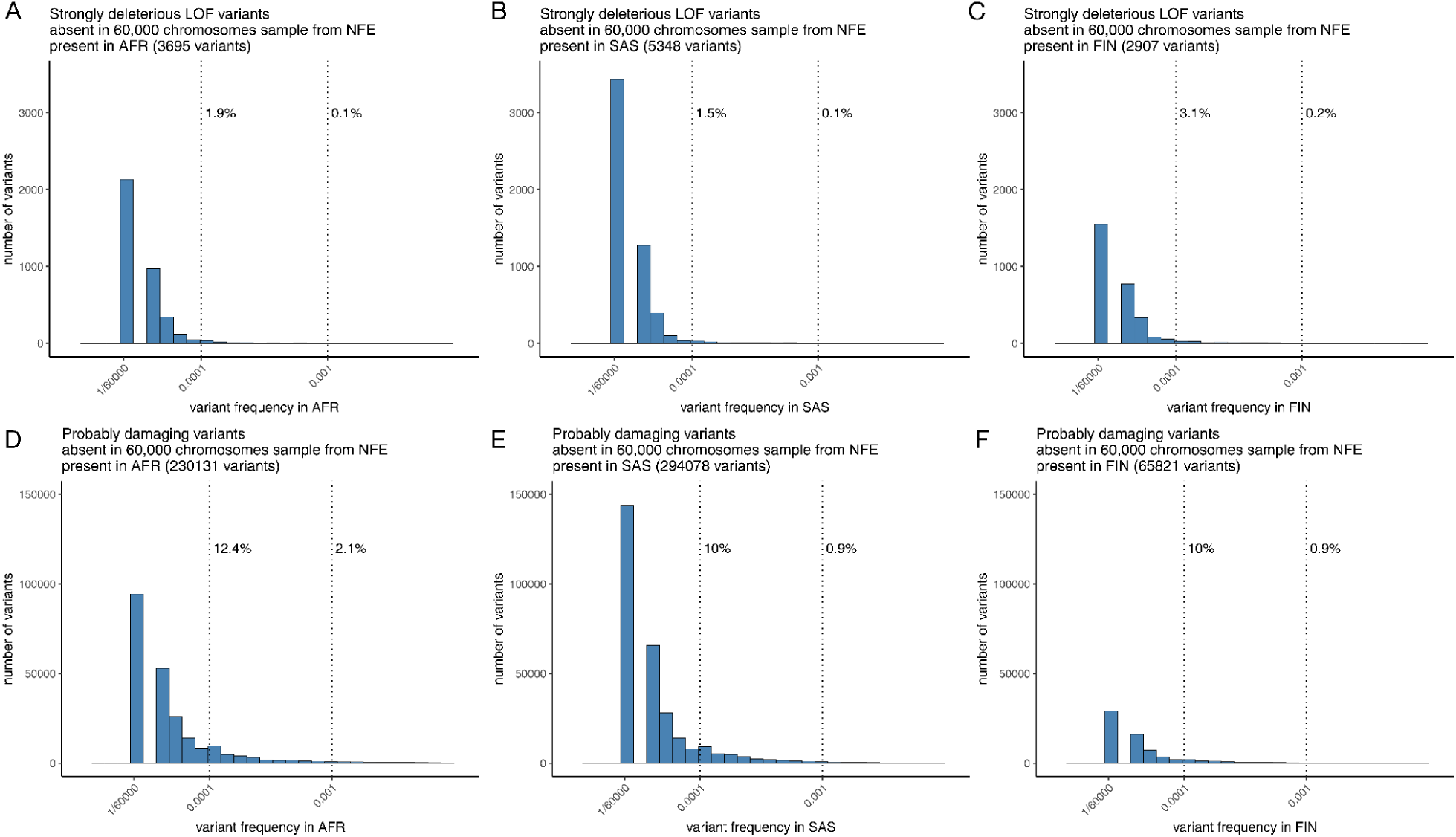
The number of variants observed in the AFR, SAS, and FIN ancestry groups, given its absence in the NFE subsample. **(A-C)** Strongly deleterious LOF variants (prior mean hs > 5%), and **(D-F)** probably damaging missense variants, as predicted by PolyPhen-2^67^. All sample sizes were downsampled to 30,000 individuals (60,000 chromosomes). Only variants absent in NFE but present in the focal ancestry group are considered. The percentages indicate the fraction of variants with a frequency >0.01% and >0.1% in the corresponding ancestry group. Unlike Supplementary Figure 6, variant frequencies here are not aggregated by gene.

**Supplementary Figure 8.**
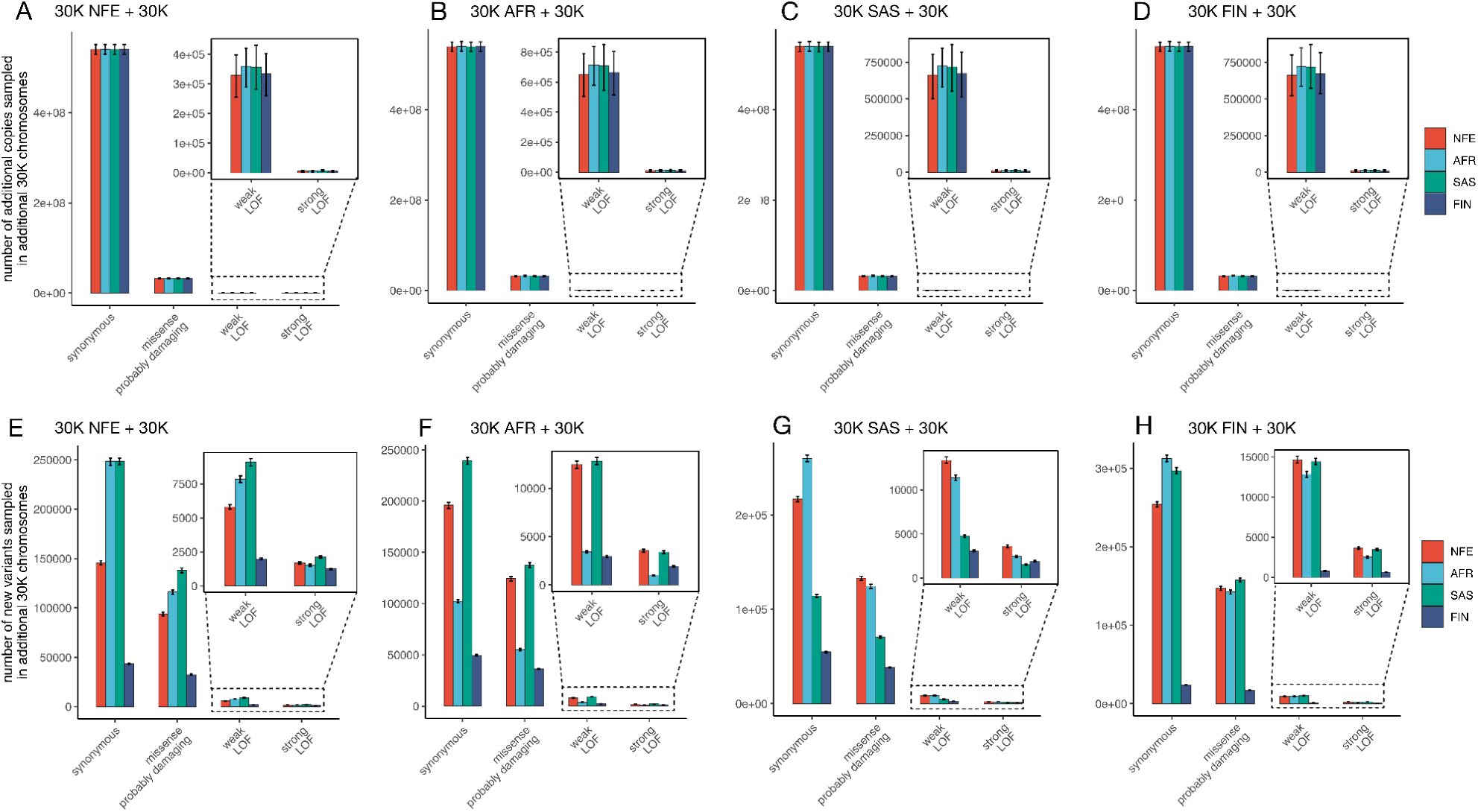
(A-D) The number of additional *copies* of derived alleles of a given functional class sampled by sequencing an additional 30,000 chromosomes from a given ancestry group in addition to 30,000 chromosomes from NFE, AFR, SAS or FIN. **(E-H)** The number of new *variants* sampled by sequencing an additional 30,000 chromosomes. The 95% confidence intervals are estimated by 10,000 bootstrap resampling of genes (see Supplementary Note 1).

**Supplementary Figure 9.**
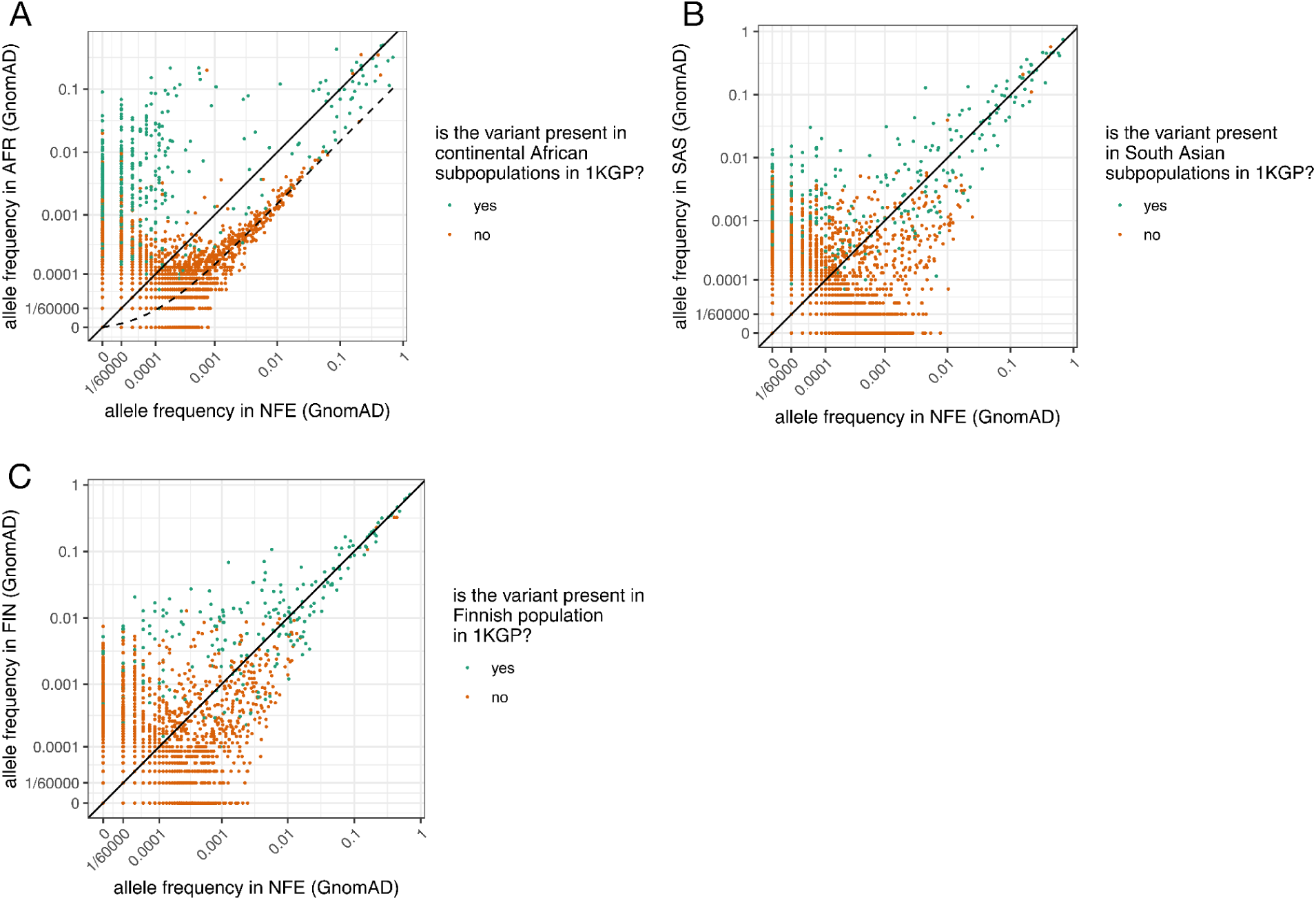
Derived allele frequencies of LOF variants in NFE versus **(A)** AFR, **(B)** SAS, or **(C)** FIN ancestry groups from gnomAD, each downsampled to 60,000 chromosomes. The dots are colored based on whether the variant is present in the corresponding non-NFE subpopulations in 1000 Genomes ^34^. For the AFR group (A), dots indicate whether the variant is present in 1000 Genomes subpopulations of African ancestry sampled in Africa, therefore excluding samples with recent European ancestry. The dashed line in (A) shows the expected allele frequency in AFR, assuming that the variant is absent in continental African populations but occurs in AFR at 15% of the frequency observed in NFE, suggesting an average of ∼15% of the recent ancestry of these samples may be shared with NFE.

## Notes

### Competing Interest Statement

The authors have declared no competing interest.

## References

1. Claussnitzer, M. et al. A brief history of human disease genetics. Nature 577, 179–189 (2020).

2. Carss, K. J. et al. Using human genetics to improve safety assessment of therapeutics. Nat Rev Drug Discov 22, 145–162 (2023).

3. Chong, J. X. et al. The Genetic Basis of Mendelian Phenotypes: Discoveries, Challenges, and Opportunities. Am J Hum Genet 97, 199–215 (2015).

4. Liggett, L. A. & Sankaran, V. G. Unraveling Hematopoiesis through the Lens of Genomics. Cell 182, 1384–1400 (2020).

5. Chakravarti, A. & Turner, T. N. Revealing rate-limiting steps in complex disease biology: The crucial importance of studying rare, extreme-phenotype families. Bioessays 38, 578–586 (2016).

6. Shendure, J. & Akey, J. M. The origins, determinants, and consequences of human mutations. Science 349, 1478–1483 (2015).

7. Agarwal, I. & Przeworski, M. Mutation saturation for fitness effects at human CpG sites. Elife 10, (2021).

8. Schraiber, J. G., Spence, J. P. & Edge, M. D. Estimation of demography and mutation rates from one million haploid genomes. bioRxiv (2024) doi:10.1101/2024.09.18.613708.

9. Mills, M. C. & Rahal, C. The GWAS Diversity Monitor tracks diversity by disease in real time. Nat Genet 52, 242–243 (2020).

10. Martin, A. R. et al. Clinical use of current polygenic risk scores may exacerbate health disparities. Nat Genet 51, 584–591 (2019).

11. Sirugo, G., Williams, S. M. & Tishkoff, S. A. The Missing Diversity in Human Genetic Studies. Cell 177, 26–31 (2019).

12. Martin, H. C. et al. Quantifying the contribution of recessive coding variation to developmental disorders. Science 362, 1161–1164 (2018).

13. Privé, F. et al. Portability of 245 polygenic scores when derived from the UK Biobank and applied to 9 ancestry groups from the same cohort. Am J Hum Genet 109, 12–23 (2022).

14. Mostafavi, H. et al. Variable prediction accuracy of polygenic scores within an ancestry group. Elife 9, (2020).

15. Hou, K. et al. Calibrated prediction intervals for polygenic scores across diverse contexts. Nat Genet 56, 1386–1396 (2024).

16. Kachuri, L. et al. Principles and methods for transferring polygenic risk scores across global populations. Nat Rev Genet 25, 8–25 (2024).

17. Wang, Y. et al. Theoretical and empirical quantification of the accuracy of polygenic scores in ancestry divergent populations. Nat Commun 11, 3865 (2020).

18. Hu, W. et al. Genomic inference of a severe human bottleneck during the Early to Middle Pleistocene transition. Science 381, 979–984 (2023).

19. The PRIMED Consortium: Reducing disparities in polygenic risk assessment. The American Journal of Human Genetics 111, 2594–2606 (2024).

20. Taliun, D. et al. Sequencing of 53,831 diverse genomes from the NHLBI TOPMed Program. Nature 590, 290–299 (2021).

21. Spence, J. P., et al. Specificity, length, and luck: How genes are prioritized by rare and common variant association studies. bioRxiv (2024) doi:10.1101/2024.12.12.628073.

22. Venner, E. et al. The frequency of pathogenic variation in the All of Us cohort reveals ancestry-driven disparities. Commun Biol 7, 174 (2024).

23. Dorschner, M. O. et al. Actionable, pathogenic incidental findings in 1,000 participants’ exomes. Am J Hum Genet 93, 631–640 (2013).

24. Wojcik, G. L. et al. Genetic analyses of diverse populations improves discovery for complex traits. Nature 570, 514–518 (2019).

25. Gurdasani, D., Barroso, I., Zeggini, E. & Sandhu, M. S. Genomics of disease risk in globally diverse populations. Nat Rev Genet 20, 520–535 (2019).

26. Sun, K. Y. et al. A deep catalogue of protein-coding variation in 983,578 individuals. Nature 631, 583–592 (2024).

27. Wang, Y. et al. Aspiring toward equitable benefits from genomic advances to individuals of ancestrally diverse backgrounds. Am J Hum Genet 111, 809–824 (2024).

28. Pubmed Search Results - ancestry-specific variants. PubMed https://pubmed.ncbi.nlm.nih.gov/?term=ancestry-specific+variants.

29. Charlesworth, B. & Charlesworth, D. Elements of Evolutionary Genetics. (Roberts & Company, Greenwood Village, CO, 2010).

30. Rands, C. M., Meader, S., Ponting, C. P. & Lunter, G. 8.2% of the Human genome is constrained: variation in rates of turnover across functional element classes in the human lineage. PLoS Genet 10, e1004525 (2014).

31. Ponting, C. P. & Hardison, R. C. What fraction of the human genome is functional? Genome Res 21, 1769–1776 (2011).

31. Ward, L. D. & Kellis, M. Evidence of abundant purifying selection in humans for recently acquired regulatory functions. Science 337, 1675–1678 (2012).

33. Lewontin, R. C. The apportionment of human diversity. in Evolutionary Biology 381–398 (Springer US, New York, NY, 1972).

34. Auton, A. et al. A global reference for human genetic variation. Nature 526, 68–74 (2015).

35. Rosenberg, N. A. et al. Genetic structure of human populations. Science 298, 2381–2385 (2002).

36. Haldane, B. S. & Innes, J. The effect of variation of fitness. American Naturalist 71, 337–349 (1937).

37. Nei, M. The frequency distribution of lethal chromosomes in finite populations. Proc Natl Acad Sci U S A 60, 517–524 (1968).

38. Crow, J. F. & Kimura, M. An Introduction to Population Genetics Theory. (New York, Evanston and London: Harper & Row, Publishers, 1970).

39. Clark, A. G. Mutation-selection balance with multiple alleles. Genetica 102–103, 41–47 (1998).

40. Fuller, Z. L., Berg, J. J., Mostafavi, H., Sella, G. & Przeworski, M. Measuring intolerance to mutation in human genetics. Nat Genet 51, 772–776 (2019).

41. Agarwal, I., Fuller, Z. L., Myers, S. R. & Przeworski, M. Relating pathogenic loss-of-function mutations in humans to their evolutionary fitness costs. Elife 12, (2023).

42. Weghorn, D. et al. Applicability of the Mutation-Selection Balance Model to Population Genetics of Heterozygous Protein-Truncating Variants in Humans. Mol Biol Evol 36, 1701–1710 (2019).

43. Zeng, T., Spence, J. P., Mostafavi, H. & Pritchard, J. K. Bayesian estimation of gene constraint from an evolutionary model with gene features. Nat Genet 56, 1632–1643 (2024).

44. Lek, M. et al. Analysis of protein-coding genetic variation in 60,706 humans. Nature 536, 285–291 (2016).

45. Karczewski, K. J. et al. The mutational constraint spectrum quantified from variation in 141,456 humans. Nature 581, 434–443 (2020).

46. Halldorsson, B. V. et al. The sequences of 150,119 genomes in the UK Biobank. Nature 607, 732–740 (2022).

47. Simons, Y. B., Turchin, M. C., Pritchard, J. K. & Sella, G. The deleterious mutation load is insensitive to recent population history. Nat Genet 46, 220–224 (2014).

48. Steiner, M. C. et al. Study design and the sampling of deleterious rare variants in biobank-scale datasets. bioRxiv 2024.12.02.626424 (2024) doi:10.1101/2024.12.02.626424.

49. Do, R. et al. No evidence that selection has been less effective at removing deleterious mutations in Europeans than in Africans. Nat Genet 47, 126–131 (2015).

50. Agrawal, A. F. & Whitlock, M. C. Inferences about the distribution of dominance drawn from yeast gene knockout data. Genetics 187, 553–566 (2011).

51. Simmons, M. J. & Crow, J. F. Mutations affecting fitness in Drosophila populations. Annu Rev Genet 11, 49–78 (1977).

52. Amorim, C. E. G. et al. The population genetics of human disease: The case of recessive, lethal mutations. PLoS Genet 13, e1006915 (2017).

53. Lachance, J. & Tishkoff, S. A. Population Genomics of Human Adaptation. Annu Rev Ecol Evol Syst 44, 123–143 (2013).

54. Quintana-Murci, L. & Barreiro, L. B. The role played by natural selection on Mendelian traits in humans. Ann N Y Acad Sci 1214, 1–17 (2010).

55. Chan, S. H. et al. Analysis of clinically relevant variants from ancestrally diverse Asian genomes. Nat Commun 13, 6694 (2022).

56. Genovese, G., Friedman, D. J. & Pollak, M. R. APOL1 variants and kidney disease in people of recent African ancestry. Nat Rev Nephrol 9, 240–244 (2013).

57. Kamiza, A. B. et al. Multi-trait discovery and fine-mapping of lipid loci in 125,000 individuals of African ancestry. Nat Commun 14, 5403 (2023).

58. Schrijver, I. et al. The Spectrum of CFTR Variants in Nonwhite Cystic Fibrosis Patients: Implications for Molecular Diagnostic Testing. J Mol Diagn 18, 39–50 (2016).

59. Coop, G. et al. The role of geography in human adaptation. PLoS Genet 5, e1000500 (2009).

60. Hernandez, R. D. et al. Classic selective sweeps were rare in recent human evolution. Science 331, 920–924 (2011).

61. Mallick, S. et al. The Simons Genome Diversity Project: 300 genomes from 142 diverse populations. Nature 538, 201–206 (2016).

62. Bhatia, G. et al. Genome-wide comparison of African-ancestry populations from CARe and other cohorts reveals signals of natural selection. Am J Hum Genet 89, 368–381 (2011).

63. Chen, S. et al. A genomic mutational constraint map using variation in 76,156 human genomes. Nature 625, 92–100 (2023).

64. Chao, K. & gnomAD Production Team. Genetic Ancestry in GnomAD v4. gnomAD browser https://gnomad.broadinstitute.org/news/2023-11-genetic-ancestry/ (2023).

65. Hudson, R. R. Genetic Data Analysis. Methods for Discrete Population Genetic Data. Bruce S. Weir. Sinauer, Sunderland, MA, 1990. xiv, 377 pp., illus. 48; paper, 27. Science 250, 575 (1990).

66. Simons, Y. B. & Sella, G. The impact of recent population history on the deleterious mutation load in humans and close evolutionary relatives. Curr Opin Genet Dev 41, 150–158 (2016).

67. Adzhubei, I., Jordan, D. M. & Sunyaev, S. R. Predicting functional effect of human missense mutations using PolyPhen-2. Curr Protoc Hum Genet Chapter 7, Unit7.20 (2013).

68. Henn, B. M., Botigué, L. R., Bustamante, C. D., Clark, A. G. & Gravel, S. Estimating the mutation load in human genomes. Nature Reviews Genetics 16, 333–343 (2015).

69. Tennessen, J. A. et al. Evolution and functional impact of rare coding variation from deep sequencing of human exomes. Science 337, 64–69 (2012).

70. An, J.-Y. et al. Genome-wide de novo risk score implicates promoter variation in autism spectrum disorder. Science 362, (2018).

71. Kaplanis, J. et al. Evidence for 28 genetic disorders discovered by combining healthcare and research data. Nature 586, 757–762 (2020).

72. Satterstrom, F. K. et al. Large-Scale Exome Sequencing Study Implicates Both Developmental and Functional Changes in the Neurobiology of Autism. Cell 180, 568–584.e23 (2020).

73. Deciphering Developmental Disorders Study. Prevalence and architecture of de novo mutations in developmental disorders. Nature 542, 433–438 (2017).

74. Blekhman, R. et al. Natural selection on genes that underlie human disease susceptibility. Curr Biol 18, 883–889 (2008).

75. Berg, J. S. et al. An informatics approach to analyzing the incidentalome. Genet Med 15, 36–44 (2013).

76. Fiziev, P. P. et al. Rare penetrant mutations confer severe risk of common diseases. Science 380, eabo1131 (2023).

77. Sham, P. C. & Purcell, S. M. Statistical power and significance testing in large-scale genetic studies. Nat Rev Genet 15, 335–346 (2014).

78. Biddanda, A., Rice, D. P. & Novembre, J. A variant-centric perspective on geographic patterns of human allele frequency variation. Elife 9, (2020).

79. Minster, R. L. et al. A thrifty variant in CREBRF strongly influences body mass index in Samoans. Nat Genet 48, 1049–1054 (2016).

80. Moltke, I. et al. A common Greenlandic TBC1D4 variant confers muscle insulin resistance and type 2 diabetes. Nature 512, 190–193 (2014).

81. Lynch, M. T. et al. The burden of pathogenic variants in clinically actionable genes in a founder population. American Journal of Medical Genetics Part A 185, 3476–3484 (2021).

82. McVean, G. A genealogical interpretation of principal components analysis. PLoS Genet 5, e1000686 (2009).

83. Gudmundsson, S. et al. Variant interpretation using population databases: Lessons from gnomAD. Hum Mutat 43, 1012–1030 (2022).

84. Jónsson, H. et al. Whole genome characterization of sequence diversity of 15,220 Icelanders. Sci Data 4, 170115 (2017).

85. Mathieson, I. & Scally, A. What is ancestry? PLoS Genet 16, e1008624 (2020).

86. Coop, G. Genetic similarity versus genetic ancestry groups as sample descriptors in human genetics. arXiv [q-bio.PE] (2022) doi:10.48550/ARXIV.2207.11595.

87. Seplyarskiy, V. et al. Cohort-level analysis of human de novo mutations points to drivers of clonal expansion in spermatogonia. medRxiv 2025.01.03.25319979 (2025) doi:10.1101/2025.01.03.25319979.

88. Kurki, M. I. et al. FinnGen provides genetic insights from a well-phenotyped isolated population. Nature 613, 508–518 (2023).

89. Kore, P. et al. Improved Allele Frequencies in gnomAD through Local Ancestry Inference. Preprint at 10.1101/2024.10.30.620961 (2024).

90. Schiffels, S. & Durbin, R. Inferring human population size and separation history from multiple genome sequences. Nat Genet 46, 919–925 (2014).

